# Genome sequence analysis of a giant-rooted ‘Sakurajima daikon’ radish (*Raphanus sativus*)

**DOI:** 10.1101/2020.02.05.936419

**Authors:** Kenta Shirasawa, Hideki Hirakawa, Nobuko Fukino, Hiroyasu Kitashiba, Sachiko Isobe

## Abstract

Daikon radish (*Raphanus sativus*) roots vary in size and shape between cultivars. This study reports the genome sequence assembly of a giant-rooted ‘Sakurajima daikon’ radish variety, ‘Okute-Sakurajima’, which produces extremely large round roots. Radish genome assembly is hampered by the repetitive and complex nature of the genome. To address this, single-molecule real-time technology was used to obtain long-read sequences at 60× genome coverage. *De novo* assembly of the long reads generated 504.5 Mb contig sequences consisting of 1,437 sequences with contig N50 length of 1.2 Mb, including 94.1% of the core eukaryotic genes. Nine pseudomolecule sequences, comprising 69.3% of the assembled contig length, were generated with high-density SNP genetic maps. The chromosome-level sequences revealed structure variations and rearrangements among Brassicaceae genomes. In total, 89,915 genes were predicted in the ‘Okute-Sakurajima’ genome, 30,033 of which were unique to the assembly in this study. The improved genome information generated in this study will not only form a new baseline resource for radish genomics, but will also provide insights into the molecular mechanisms underlying formation of giant radish roots.

## 1. Introduction

Daikon radish (*Raphanus sativus*) is a member of the Brassicaceae family of flowering plants. Daikon radish roots of different cultivars vary considerably in their size and shape.^1^ For example, among daikon cultivars, ‘Sakurajima daikon’ radishes exhibit the largest roots. ‘Sakurajima daikon’ is the name of a group of daikon radish varieties that are mainly cultivated in the Kagoshima prefecture of Japan. Soil in this region contains volcanic ash from Mt. Sakurajima and is thought to be particularly suitable for cultivation of ‘Sakurajima daikon’. One well-known ‘Sakurajima daikon’ line is ‘Okute-Sakurajima’, which has large round roots that can exceed 20 kg.^1^ The size and shape of ‘Okute-Sakurajima’ roots are desirable traits for plant breeding, but the molecular mechanisms underpinning these characteristics remain unknown.

Four genome assemblies based on next-generation sequencing technologies have been reported for radish,^2–5^ but these do not include ‘Okute-Sakurajima’ or ‘Sakurajima daikon’. The sequence contiguities of the genome assemblies are relatively short, and the entire radish genome is not covered.^6^ One reason for the short assembly size may be the complexity of the radish genome because *Raphanus*, like other species in the *Brassica* genus, underwent genome triplication sometime after divergence from Arabidopsis.^7^ Furthermore, radish, in common with other *Brassica*, is self-incompatible and allogamous, resulting in a highly heterozygous genome.^8^

Recent advances in long-read sequence technology have allowed highly heterozygous genomes of several plant species to be successfully sequenced.^9^ In this study, the ‘Okute-Sakurajima’ genome was sequenced using long-read technology. Sequences were aligned to radish chromosomes, establishing pseudomolecules and allowing gaps in previous radish genome assemblies to be resolved. This enhanced radish genome will provide insights into radish evolutionary development and, specifically, into the molecular underpinnings of giant daikon root formation.

## 2. Materials and methods

### 2.1. ‘Okute-Sakurajima’ *de novo* genome assembly

Total DNA was extracted from young leaves of the ‘Okute-Sakurajima’ daikon radish cultivar (NARO GeneBank accession number: JP27228) using a Genomic-tip kit (Qiagen, Hilden, Germany). Short-read sequence data were obtained using a MIGSEQ-2000 DNA sequencer (also known as a DNBSEQ-G400; MGI Tech, Shenzhen, China) and were used to estimate the size of the ‘Okute-Sakurajima’ genome, with *k*-mer distribution analysis performed using ‘Jellyfish’. To gain long-read sequence data, an SMRT sequence library was constructed with an SMRTbell Express Template Prep Kit (PacBio, Menlo Park, CA, USA) and sequenced on a PacBio Sequel system (PacBio). The long reads from the Sequel system were assembled, and the haplotypes were phased with ‘Falcon-unzip’. The assembly was polished twice with ‘Arrow’ and designated as RSAskr_r1.0. Assembly completeness was evaluated with ‘BUSCO’.

### 2.2. Construction of map-based pseudomolecule sequences

An F2 mapping population (n=115), termed SNF2, derived from a cross between an inbred line via self-pollination of radish cultivar ‘Shogoin Daikon’ and a line of *R. sativus* var. *raphanistroides* ‘Nohara 1’, collected from Nohara, Maizuru, Kyoto, Japan, was used to establish genetic maps according to the methods of Shirasawa and Kitashiba.^6^ In brief, DNA was extracted from leaves of each line and used for ddRAD-Seq library construction. The library was sequenced on a HiSeq4000 sequencer (Illumina, San Diego, CA, USA). Data analysis was also performed as described by Shirasawa and Kitashiba.^6^ After trimming low-quality sequences and adapter sequences using ‘FASTX-Toolkit’ and ‘PRINSEQ’, the remaining high-quality reads were mapped onto the RSAskr_r1.0 assembly, using ‘Bowtie2’ to call SNPs using the mpileup command in ‘SAMtools’ followed by filtering out the low-quality data with ‘VCFtools’. In parallel, ddRAD-Seq data (DRA accession number: DRA005069) from another F2 population (n=95), namely ASF2,^2^ derived from a cross between ‘Aokubi *S-h*’ and ‘Sayatori 26704’, was also analyzed as above. SNP data were used for genetic map construction with ‘Lep-Map3’. On the basis of genetic maps constructed with ‘ALLMAPS’, the RSAskr_r1.0 sequence assembly was assigned to radish chromosomes to produce pseudomolecule sequences, termed RSAskr_r1.0.pmol. The genome structure of the ‘Okute-Sakurajima’ genome was compared with those of *R. sativus*,^4^ *Brassica rapa*,^10^ and *Arabidopsis thaliana*^11^ using ‘D-Genies’.

### 2.3. Gene identification in the genome sequences

The gene models predicted in the RSA_r1.0 assembly^2^ were mapped onto the RSAskr_r1.0.pmol pseudomolecule sequences using ‘Minimap2’. In addition, *ab initio* gene prediction was performed on the RSAskr_r1.0.pmol sequences using ‘Augustus’ as described in Kitashiba et al.^2^

## 3. Results

### 3.1. *De novo* assembly of the ‘Okute-Sakurajima’ radish genome

*K*-mer distribution analysis of the 147.3 Gb short-read data indicated that the ‘Okute-Sakurajima’ radish genome was highly heterozygous, and that the estimated haploid genome size was 592.4 Mb (Supplementary Figure S1). Subsequent long-read sequencing produced 36.0 Gb data (60.7× coverage of the estimated genome size) with 2.3 million reads with N50 length of 29.1 kb. After two rounds of polishing, the long-read assembly consisted of 504.5 Mb primary contigs (including 1,437 sequences with N50 length of 1.2 Mb) and 263.5 Mb haplotig sequences (including 2,373 sequences with N50 length of 154.6 kb) (Table 1). A BUSCO analysis of the primary contigs indicated that 71.0% and 23.1% of sequences were single-copy complete BUSCOs and duplicated complete BUSCOs, respectively (Table 1), suggesting that most of the gene regions were represented in the primary contigs.

**Table 1.** Statistics of the primary contig sequences of ‘Okute-Sakurajima’

|  | RSAskr_r1.0 |
| --- | --- |
| Total contig size (bases) | 504,534,164 |
| Number of contigs | 1,437 |
| N50 length (bases) | 1,247,688 |
| Longest contig size (bases) | 8,317,732 |
| Gap (%) | 0.0 |
| Complete BUSCOs | 94.1 |
| (Single-copy BUSCOs | 71.0) |
| (Duplicated BUSCOs | 23.1) |
| Fragmented BUSCOs | 2.0 |
| Missing BUSCOs | 3.9 |
| #Genes (*ab initio*) | 89,915 |
| #Genes (mapping) | 78,645 |

### 3.2. Construction of pseudomolecule sequences based on genetic maps

In total, 5,872 and 2,830 high-quality SNPs were obtained from the SNF2 and ASF2 mapping populations, respectively, and employed for linkage analysis. The resultant genetic map for SNF2 consisted of nine linkage groups with 5,570 SNPs covering 867.2 cM, and the map for ASF2 comprised nine linkage groups with 2,680 SNPs covering 895.3 cM (Supplementary Tables S2). Contig sequences of RSAskr_r1.0 were assigned to the radish chromosomes in accordance with the two genetic maps. Nine pseudomolecule sequences, termed RSAskr_r1.0.pmol, spanning 349.8 Mb (69.3%) were established with 293 contigs (Table 2), of which 95 sequences (218.5 Mb, 43.3%) were oriented. The nine resulting sequences were named using the nomenclature (R1–R9) proposed by Shirasawa and Kitashiba^6^. The remaining unassigned sequences (n=1,144, 154.7 Mb, 30.7%) were designated as R0. The structure of the ‘Okute-Sakurajima’ genome was conserved in *R. sativus* but partially disrupted in *B. rapa* and *A. thaliana*, as indicated previously^2^.

**Table 2.** Statistics of the ‘Okute-Sakurajima’ pseudomolecule sequences, RSAskr_r1.0.pmol.

| Chr | #Contigs | (%) | Contig size (bp) | (%) | #Genes | (%) |
| --- | --- | --- | --- | --- | --- | --- |
| R1 | 25 | 1.7 | 27,719,058 | 5.5 | 5,426 | 6.0 |
| R2 | 40 | 2.8 | 42,944,316 | 8.5 | 8,295 | 9.2 |
| R3 | 30 | 2.1 | 31,410,669 | 6.2 | 5,923 | 6.6 |
| R4 | 35 | 2.4 | 56,498,296 | 11.2 | 10,843 | 12.1 |
| R5 | 34 | 2.4 | 42,357,306 | 8.4 | 8,477 | 9.4 |
| R6 | 41 | 2.9 | 53,940,652 | 10.7 | 10,403 | 11.6 |
| R7 | 17 | 1.2 | 28,108,325 | 5.6 | 5,545 | 6.2 |
| R8 | 30 | 2.1 | 28,319,830 | 5.6 | 5,474 | 6.1 |
| R9 | 41 | 2.9 | 38,520,653 | 7.6 | 7,143 | 7.9 |
| **R1-R9** | **293** | **20.4** | **349,819,105** | **69.3** | **67,529** | **75.1** |
| R0 | 1,144 | 79.6 | 154,715,059 | 30.7 | 22,386 | 24.9 |
| **Total** | **1,437** | **100.0** | **504,534,164** | **100.0** | **89,915** | **100.0** |

### 3.3. Genes on the ‘Okute-Sakurajima’ genome sequence

In total, 89,915 gene models were predicted in the RSAskr_r1.0 assembly using an *ab initio* gene prediction method (Table 2). To assess the availability of gene spaces in RSAskr_r1.0,^2^ the 80,521 predicted gene models were aligned to the RSAskr_r1.0.pmol pseudomolecules. Of the predicted gene models, 78,645 (97.6%) were aligned, suggesting that most of the genes were represented in the RSAskr_r1.0.pmol assembly. The genome positions of 59,882 predicted genes of RSAskr_r1.0 and 77,496 mapped genes of RSA_r1.0 overlapped. The remaining 30,033 genes (=89,915-59,882) were unique to the assembled sequences of ‘Okute-Sakurajima’.

## 4. Discussion

In this study, we report the genome sequence assembly of ‘Okute-Sakurajima’ radish, a variety of ‘Sakurajima-diakon’, based on long-read sequence technology. The total assembly size of 504.5 Mb is the longest reported for any radish genome to date,^2–5^ suggesting that the long reads might span the repetitive sequences found throughout the radish genome. However, since the assembly size was approximately 90 Mb shorter than the estimated genome size, the long sequencing technology was not sufficient to fully resolve the difficulties in assembling the complex genome of this mesopolyploid species.^7^

Map-based pseudomolecule sequences comprising 69.3% of assembled sequences were produced. Unexpectedly, 30.7% of assembled sequence remained unassigned to radish chromosomes. As genetic mapping is reliant upon SNP availability in the genome, sequences cannot be assigned where SNPs are not present. To overcome this genetic limitation, optical mapping and Hi-C technologies, both of which are based on physical mapping strategies, have been developed.^9^ These technologies, alongside traditional genetic mapping, would allow the genome coverage of the assembly and the completeness of pseudomolecules to be further improved.

In this study, 30,033 genes were discovered that were unique to the current assembly, suggesting that these genes may not have been identified in previous studies.^2^ The expanded genomic information obtained in this study is expected to provide new insights into radish growth and development. For example, this study is the first genome report for a giant radish cultivar, and some of the genes unique to the ‘Okute-Sakurajima’ genome may be involved in giant root formation. Further comparative analysis with radish cultivars with divergent root shapes and sizes would provide insights into the genetic mechanisms contributing to giant root formation.

**Figure 1.**
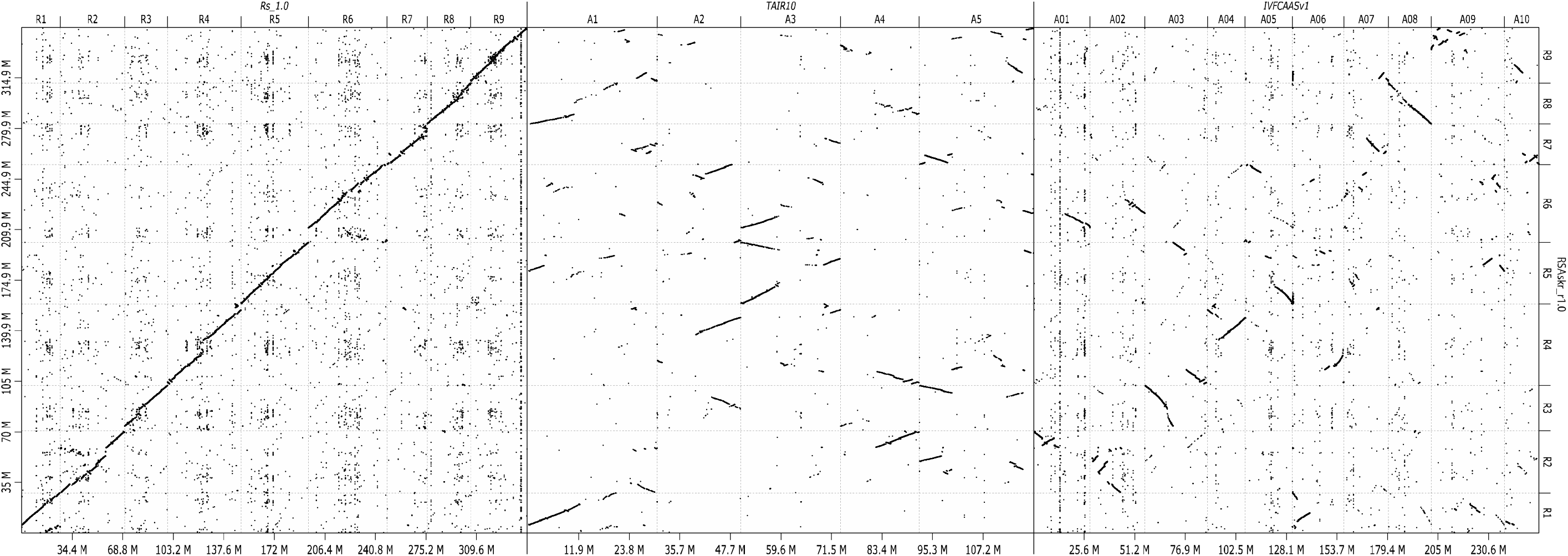
Comparative maps of the ‘Okute-Sakurajima’ genome. Dots indicate sequence similarities of the ‘Okute-Sakurajima’ genome (RSAskr_r1.0) on the vertical axis with those of *R. sativus* (Rs_1.0), *A. thaliana* (TAIR10), and *B. rapa* (IVFCAASv1) on the horizontal axis.

## Footnote

References for data analysis tools used in this study, which are indicated with back quotes in the text, are listed in Supplementary Table S1.

## Acknowledgments

We thank S. Sasamoto, S. Nakayama, A. Watanabe, T. Fujishiro, Y. Kishida, M. Kohara, C. Minami, H. Tsuruoka, and M. Yamada (Kazusa DNA Research Institute) for their technical assistance. ‘Okute-Sakurajima’ seeds (Accession number: JP27228) were provided by the NARO GeneBank, Tsukuba, Japan.

## Data availability

Sequence reads are available from the DNA Data Bank of Japan (DDBJ) Sequence Read Archive (DRA) under the accession number DRA009553. The DDBJ accession numbers of assembled sequences are BLLE01000001-BLLE01003810. Genome information is available at Plant GARDEN (https://plantgarden.jp).

## Funding

This work was supported by the Kazusa DNA Research Institute Foundation and the Project of the NARO Bio-oriented Technology Research Advancement Institution (Research program on development of innovative technology, Grant number: 29010B).

## Conflict of interest

None declared.

## Supplementary Data

**Supplementary Table S1** Program tools used for genome assembly and gene prediction.

**Supplementary Table S2** Genetic map length and number of SNPs for F2 radish populations.

**Supplementary Figure S1** Genome size estimation for ‘Okute-Sakurajima’ with the distribution of the number of distinct *k*-mers (*k*=17) with the given multiplicity values.

## References

1. Yamagishi, H. 2017, Speciation and Diversification of Radish. In: Nishio, T. and Kitashiba, H. (eds), The Radish Genome, Springer, Cham, pp. 11–30.

2. Kitashiba, H., Li, F., Hirakawa, H., et al. 2014, Draft sequences of the radish (*Raphanus sativus* L.) genome. DNA Res, 21, 481–490.

3. Mitsui, Y., Shimomura, M., Komatsu, K., et al. 2015, The radish genome and comprehensive gene expression profile of tuberous root formation and development. Sci Rep, 5, 10835.

4. Jeong, Y. M., Kim, N., Ahn, B. O., et al. 2016, Elucidating the triplicated ancestral genome structure of radish based on chromosome-level comparison with the *Brassica* genomes. Theor Appl Genet, 129, 1357–1372.

5. Zhang, X., Yue, Z., Mei, S., et al. 2015, A *de novo* genome of a Chinese radish cultivar. Hort Plant J, 1, 155–164.

6. Shirasawa, K. and Kitashiba, H. 2017, Genetic Maps and Whole Genome Sequences of Radish. In: Nishio, T. and Kitashiba, H. (eds), The Radish Genome, Springer, Cham, pp. 31–42.

7. Moghe, G. D., Hufnagel, D. E., Tang, H., et al. 2014, Consequences of Whole-Genome Triplication as Revealed by Comparative Genomic Analyses of the Wild Radish Raphanus raphanistrum and Three Other Brassicaceae Species. Plant Cell, 26, 1925–1937.

8. Nishio, T. and Sakamoto, K. 2017, Polymorphism of Self-Incompatibility Genes. In: Nishio, T. and Kitashiba, H. (eds), The Radish Genome, Springer, Cham, pp. 177–188.

9. Michael, T. P. and VanBuren, R. 2020, Building near-complete plant genomes. Curr Opin Plant Biol, 54, 26–33.

10. Wang, X., Wang, H., Wang, J., et al. 2011, The genome of the mesopolyploid crop species *Brassica rapa*. Nat Genet, 43, 1035–1039.

11. Arabidopsis Genome, I. 2000, Analysis of the genome sequence of the flowering plant *Arabidopsis thaliana*. Nature, 408, 796–815.

