## Supplementary Figure 1 for "Genome sequence analysis of a giant-rooted ‘Sakurajima daikon’ radish (*Raphanus sativus*)"

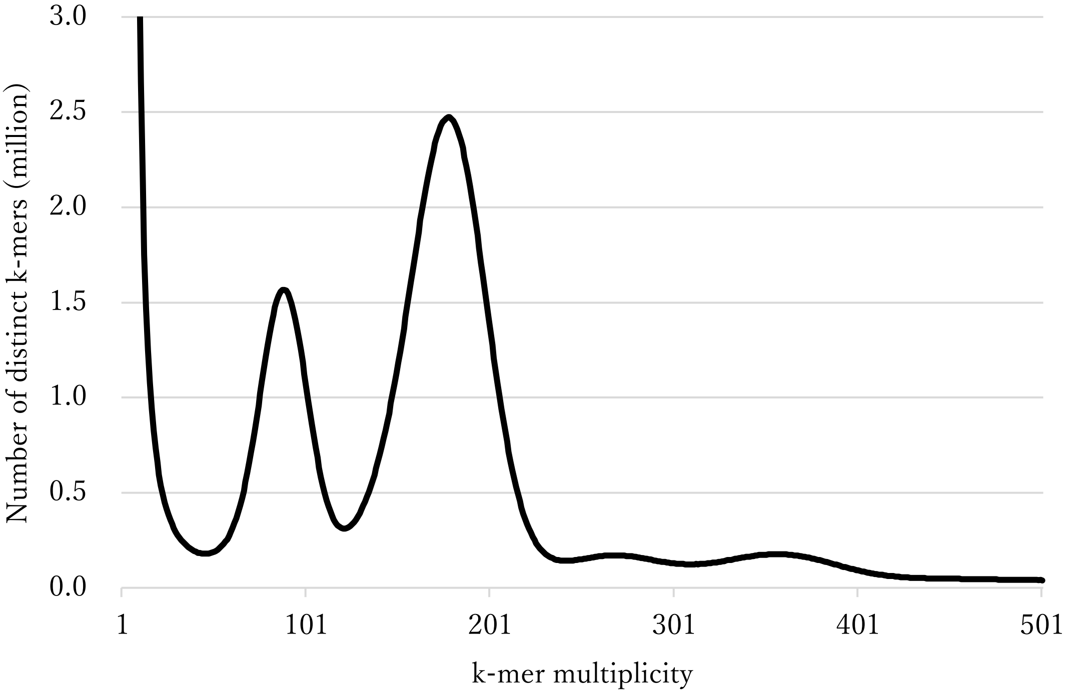


**Supplementary Figure S1.** Genome size estimation for 'Okute-Sakurajima' with the distribution of the number of distinct *k*-mers (*k*=17) with the given multiplicity values.
